# Guidance for high-quality functional gene embeddings from large language models

**DOI:** 10.64898/2026.04.30.721875

**Authors:** Rongyao Huang, Yaopan Hou, Wuye Zhao, Junbing Zhang, Jian Lu, Yimeng Kong, Peng Xu

## Abstract

Large language models (LLMs) are increasingly used to generate gene embeddings, yet systematic benchmarks of prompting strategies and practical guidance for obtaining biologically meaningful representations remain limited. Here we present GEbench, an evaluation framework for assessing LLM-derived gene embeddings across different tasks, prompting strategies, and LLM architectures. GEbench revealed that embedding quality depends primarily on whether the input text contains explicit functional information, rather than on sparse gene identifiers or model size. Identifier-based embeddings showed weak biological organization, whereas embeddings derived from functional descriptions consistently achieved stronger functional separation and predictive performance. Notably, Self-Des, which extracts embeddings from model-generated gene function descriptions, enabled locally deployable LLMs to generate high-fidelity representations that approach the quality of expert-curated databases. Genome-scale analyses further supported these findings, indicating that explicit functional descriptions are an effective design principle for generating high-quality gene embeddings from LLMs.

## Main

Large language models (LLMs) are emerging as a promising source of gene representations for downstream genomics analyses, including pathway inference, network modeling, and functional prediction^1-4^. Unlike embeddings derived from curated annotations or task-specific neural architectures, LLM-based embeddings can convert textual biological knowledge into a shared representation space with broad applicability. However, current studies vary substantially in model architecture, input formulation, and evaluation strategy, making it difficult to determine which inputs produce embeddings that best reflect gene function^5-7^. In practice, guidance on obtaining high-quality functional gene embeddings from LLMs remains limited.

A central limitation is the lack of a standardized, biologically grounded evaluation framework. Many existing assessments rely on isolated downstream tasks or ad hoc metrics, which can obscure the factors that determine embedding quality and complicate comparisons across models and prompting strategies. In particular, current studies often do not distinguish whether improved performance arises from richer biological information in the input text or from model-specific technical choices. To address this gap, we developed GEbench, a compact benchmark designed to evaluate whether gene embeddings preserve biologically meaningful functional structure.

GEbench integrates three complementary tasks: unsupervised gene clustering, supervised pathway prediction, and protein–protein interaction (PPI) prediction (**Fig. 1a**). To construct a stringent functional reference space, we integrated pathway and ontology annotations from KEGG^8^, Reactome^9^, and Gene Ontology^10^, and generated a binary gene–function association matrix followed by iterative dimensionality reduction and unsupervised clustering (**Methods**). This procedure yielded a compact ground-truth set of 680 human genes distributed across five coherent, well-separated functional modules representing canonical biological programs: allograft rejection, apical junction, E2F targets, fatty acid metabolism, and TGF-β signaling (Supplementary **Fig. S1**). These modules served as a common reference for comparing embedding strategies across models.

**Fig. 1.**
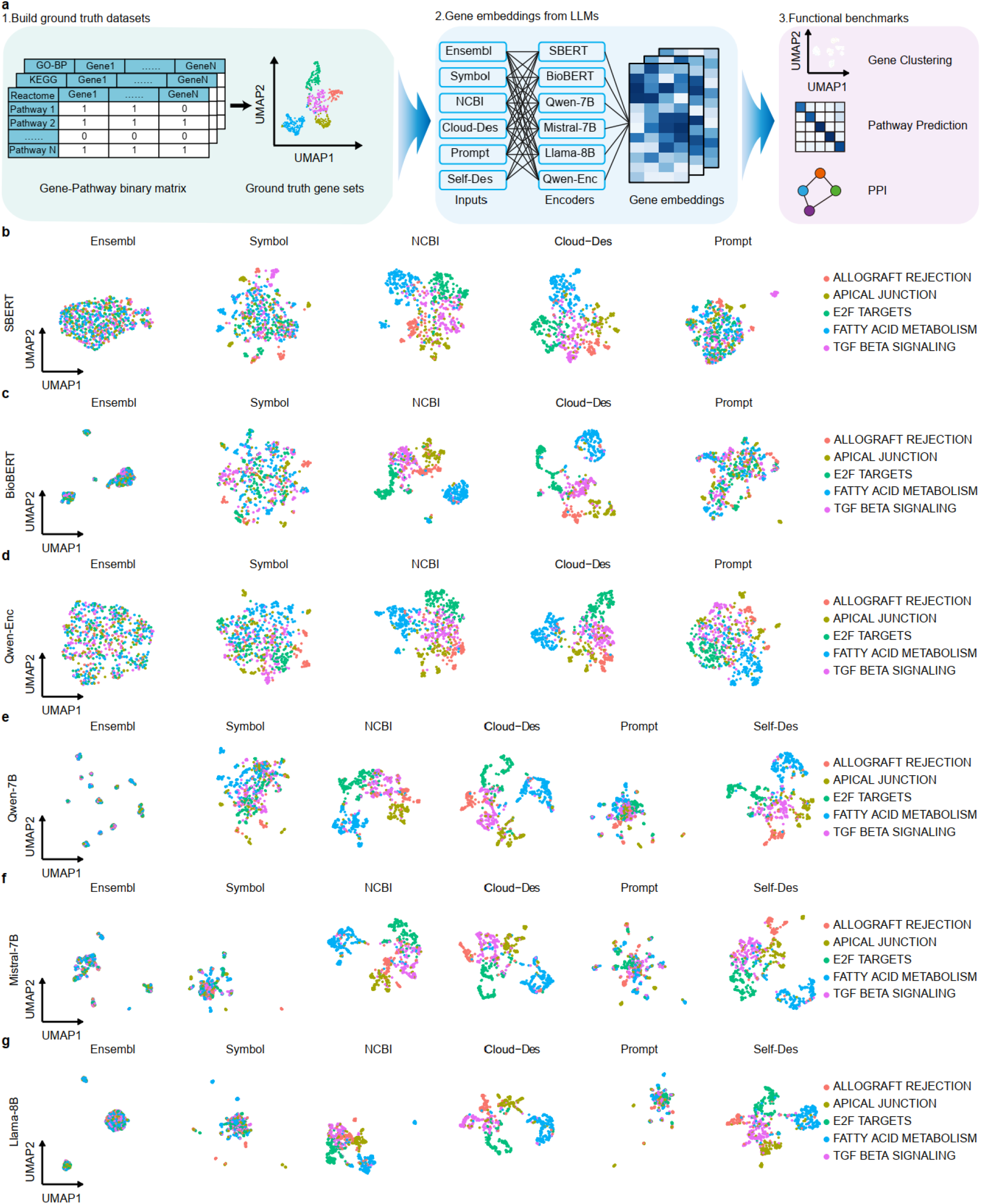
GEbench framework and functional organization of gene embeddings generated by language models. **a**, Overview of the GEbench framework. A curated functional reference space was constructed from pathway and ontology databases and used to evaluate gene embeddings across three complementary tasks: unsupervised gene clustering, supervised pathway prediction, and protein-protein interaction prediction. Embeddings were generated from six input types: Ensembl, Symbol, NCBI, Cloud-Des, Prompt, and Self-Des. **b–g**, UMAP projections of gene embeddings generated by representative encoder- and decoder-based models, including SBERT (**b**), BioBERT (**c**), Qwen-Enc (**d**), Qwen-7B (**e**), Mistral-7B (**f**) and Llama-8B (**g**). Colors indicate benchmark ground-truth functional groups. Embeddings derived from explicit functional descriptions, particularly Self-Des, form compact and well-separated clusters, whereas identifier-based embeddings remain diffuse and overlapping.

## Results

We applied GEbench to embeddings generated by both encoder- and decoder-based language models. Encoder models included SBERT^11^, BioBERT^12^, and the cloud-based Qwen-Enc embedding service. Decoder models contained locally deployed models, including Qwen-7B, Mistral-7B, and Llama-8B. To determine how input formulation affects functional representation quality, we compared six input strategies: Ensembl, Symbol, NCBI, Cloud-Des, Prompt, and Self-Des. Ensembl and Symbol represent sparse identifier inputs. NCBI corresponds to expert-curated gene summaries. Cloud-Des denotes functional descriptions generated by a cloud-based full-parameter LLM service. Prompt refers to embeddings extracted directly from the full instruction prompt containing the gene symbol, whereas Self-Des denotes embeddings extracted from a functional description generated by the same local model. This design allowed us to distinguish the effects of sparse identifiers, external descriptive text, and self-generated functional descriptions within a unified evaluation framework.

Embeddings derived from sparse identifiers showed weak functional organization across all tested models. In UMAP space, Ensembl- and Symbol-based embeddings remained diffuse and overlapping, with poor separation of the benchmark functional groups (**Fig. 1b–g**). This pattern was confirmed quantitatively by adjusted Rand index (ARI), where identifier-based embeddings performed near random expectation in the clustering benchmark (**Fig. 2a**). Thus, sparse identifiers alone were generally insufficient for generating functionally informative gene embeddings. Although gene symbols may trigger latent biological knowledge in LLMs, this knowledge was not consistently encoded in a representation space that preserved functional structure.

**Fig. 2.**
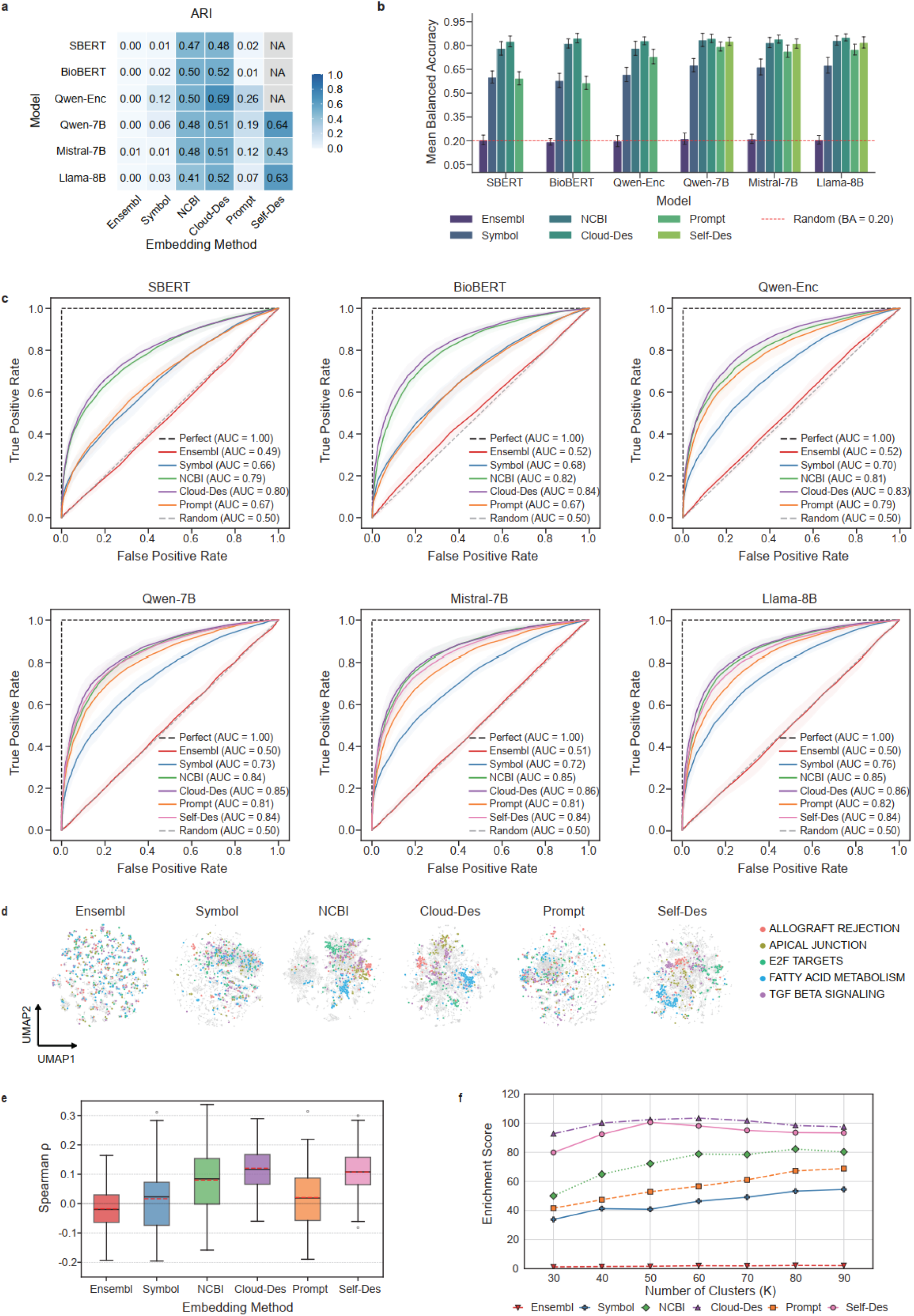
Multi-task evaluation of functional gene embeddings using GEbench. **a**, Clustering performance measured by adjusted Rand index (ARI), comparing k-means assignments with benchmark ground-truth labels. Inputs containing explicit functional descriptions consistently recover functional structure more accurately than identifier-based or prompt-based inputs. **b**, Pathway prediction performance measured by balanced accuracy of a five-class random forest classifier trained on gene embeddings. Error bars indicate standard error across repeated cross-validation. **c**, Protein–protein interaction prediction performance evaluated by ROC curves and mean AUROC values. Description-based inputs consistently outperform identifier-based embeddings. **d**, UMAP visualization of genome-scale embeddings generated from different input strategies. Grey points represent all protein-coding genes, and colored points represent genes from representative biological pathways. **e**, Correlation between pairwise embedding similarity and Gene Ontology Biological Process similarity. Inputs containing explicit functional descriptions show the strongest agreement with ontology-based functional relatedness. **f**, Functional coherence of genome-scale embeddings measured by clustering followed by Gene Ontology enrichment analysis across a range of cluster numbers. Description-based inputs consistently achieve higher functional enrichment.

In contrast, embeddings derived from text containing explicit functional descriptions showed substantially stronger biological organization. Both NCBI and Cloud-Des inputs yielded compact, well-separated clusters corresponding to known biological programs (**Fig. 1b–g**). This advantage extended across all three evaluation tasks. In pathway prediction, description-based inputs consistently achieved higher balanced accuracy than identifier-based inputs (**Fig. 2b**). In PPI prediction, embeddings derived from explicit functional descriptions also showed markedly improved performance, with AUROC values reaching 0.80–0.86 across decoder-based models (**Fig. 2c**). Across models and tasks, the strongest determinant of embedding quality was therefore not the identifier type or model scale, but whether the input text explicitly described gene function.

We next asked whether a practical local strategy in decoder-based models (Qwen-7B, Mistral-7B, and Llama-8B) could reproduce this advantage without relying on external curated resources. In these locally deployed models, Prompt modestly improved performance relative to Symbol, indicating that the inclusion of functional instruction text can partially enrich representation quality. However, Self-Des consistently outperformed Prompt across all evaluated decoder models. In the clustering task, Self-Des achieved markedly higher ARI values and approached the performance obtained with NCBI and Cloud-Des (**Fig. 2a**). Similar gains were observed in pathway prediction and PPI prediction (**Fig. 2b**,**c**). These results indicate that local decoder-based models benefit strongly when embeddings are derived from a stand-alone gene-function statement rather than from prompt text alone.

To determine whether this guidance generalized beyond the benchmark gene set, we extended the analysis to 18,954 human protein-coding genes. Genome-scale embeddings derived from functional descriptions showed stronger agreement with Gene Ontology Biological Process similarity and higher cluster-level functional coherence than embeddings derived from Ensembl, Symbol, or Prompt inputs (**Fig. 2d–f**). The relative ordering of input strategies was consistent with the benchmark results, indicating that the conclusions derived from GEbench generalized to a genome-wide setting.

## Discussion

Our results provide a clear practical message for generating functional gene embeddings from LLMs. Sparse identifiers such as Ensembl IDs and gene symbols should not be used alone when the goal is to obtain biologically meaningful functional representations. Although LLMs may contain latent knowledge linked to these identifiers, this knowledge is not consistently encoded or retrieved in an embedding space that preserves functional structure. By contrast, inputs containing explicit functional descriptions consistently yield stronger biological organization and better predictive performance across models and tasks.

The comparison between Prompt and Self-Des further clarifies the source of this improvement. Prompt embeddings are extracted from input text that still contains the instructional frame together with the gene symbol. Self-Des, instead, uses a direct functional description generated by the model itself as the embedding input. The superior performance of Self-Des therefore suggests that the benefit does not arise simply from increasing text length, but from using text with clearer biological semantics. In practical terms, transforming sparse identifiers into standalone functional statements emerges as a simple yet effective local strategy for improving decoder-based gene embeddings.

Together, these analyses provide practical guidance for generating high-quality functional gene embeddings from LLMs. Across models and tasks, embedding quality depended primarily on whether the input text contained explicit functional information, rather than on sparse gene identifiers or even model size. For locally deployable decoder-based models, Self-Des offers a simple yet effective strategy to improve embedding quality. More broadly, our findings establish explicit functional description as a key design principle for biologically meaningful representation learning from LLMs.

## Methods

### Model selection and deployment

Six language models were selected for embedding evaluation, representing both encoder- and decoder-based architectures. Encoder models included Sentence-BERT (SBERT), BioBERT, and the cloud-based full-parameter Qwen text-embedding-v4 (Qwen-Enc). Decoder models included Qwen2.5-7B-Instruct (Qwen-7B), Mistral-7B-Instruct-v0.3 (Mistral-7B), and Llama-3.1-8B-Instruct (Llama-8B). Additionally, the cloud-based Qwen-Turbo model was employed specifically for generating semantic descriptions of gene functions (Qwen-Des). Embedding dimensions varied by model architecture: SBERT (384), BioBERT (1,024), Qwen-7B (3,584), and both Mistral-7B and Llama-8B (4,096). For Qwen-Enc, we utilized the default dimension of 1,024. Regarding deployment, SBERT, BioBERT, Qwen-7B, Mistral-7B, and Llama-8B were obtained from ModelScope (https://www.modelscope.cn/models) and deployed on a local server equipped with an Apple M4 chip (Mac Studio). Local inference was accelerated using the Metal Performance Shaders (MPS) backend to ensure computational efficiency and reproducibility. The Qwen-Enc and Qwen-Turbo models were accessed via the Alibaba Cloud DashScope API.

To ensure deterministic and reproducible text generation, a standardized parameter configuration was applied for both D-Prompt and Qwen-Des descriptions. The temperature parameter was set to 0.0 to maximize output consistency, and a fixed seed value of 42 was utilized to eliminate stochastic variability. The maximum token limit was set to 256 to constrain response length. Manual inspection confirmed that outputs under these settings were conceptually stable across repetitions.

### Software environment and embedding extraction

Gene embeddings were extracted via a standardized computational pipeline implemented in Python 3.10.16, leveraging transformers (v4.45.3) and torch (v2.6.0). The extraction protocol was executed according to the following four technical specifications:

1. Precision control: Models were loaded with bfloat16 precision. The configuration was adopted to preserve a high numerical dynamic range for latent representations while optimizing memory efficiency on the unified memory architecture.
2. Layer selection and feature capture: During the forward pass, the output_hidden_states parameter was enabled to access the model’s internal representations. We specifically extracted the final hidden state, as it provides the most semantically integrated representation for downstream analysis.
3. Aggregation via masked mean pooling: To condense variable-length sequence information into a singular fixed-length vector, we applied masked mean pooling. The process calculated the arithmetic mean across all valid token embeddings, explicitly excluding padding tokens via the attention_mask.
4. Numerical stabilization: To ensure computational stability during batch processing, a clamping factor of 1 × 10^™9^ was applied to the mask summation. This safeguard prevented division-by-zero errors during the averaging step, ensuring consistent embedding normalization.

### Prompting strategies and input formulations

Human protein-coding genes were acquired from the Ensembl database (GRCh38.p14). The biomaRt package^13^ was used to convert gene symbols to Ensembl IDs. For symbols with multiple mappings, a single identifier was randomly selected. We filtered the dataset to exclude genes lacking a corresponding gene symbol, yielding 19,477 protein-coding genes. Subsequently, we queried the NCBI Gene database and retrieved gene summaries for 18,954 of these genes. To ensure consistency across all analyses, the final study cohort was restricted to these 18,954 genes for which both Ensembl IDs and NCBI gene summaries were available. To evaluate the impact of input modality on embedding quality, we designed six distinct prompting strategies categorized into identifier-based, external-knowledge-based, and LLM-generated inputs:

1. Identifier-based inputs (Ensembl ID and Gene Symbol): To establish a baseline, we generated embeddings using raw identifiers. For Ensembl IDs, unique gene identifiers (e.g., *ENSG00000141510*) were used as direct inputs. For Gene Symbols, the standard HGNC symbol (e.g., *TP53*) was used as the sole input token.
2. External knowledge input (NCBI): To assess embeddings derived from expert-curated text, we retrieved full gene summaries from the NCBI Gene database. The complete abstract text served as the input for the embedding models.
3. Cloud-generated description (Cloud-Des): To evaluate the utility of high-quality synthetic text, we generated functional descriptions using the full-parameter Qwen-Turbo model via the Alibaba Cloud API. We employed a standardized instruction: “Describe biological functions of the [Gene Symbol] in one sentence.” The resulting generated text was then used as the input for the embedding models.
4. Prompt input: This strategy evaluates the model’s internal knowledge without a distinct generation step. The model is provided with the context instruction “Describe biological functions of the [Gene Symbol] in one sentence,” and the embedding is extracted directly from this prompt context.
5. Self-generated functional descriptions (Self-Des): We developed a two-stage strategy to decouple semantic generation from representation learning. In the first stage, the locally deployed LLM (Qwen-7B, Mistral-7B, or Llama-8B) generates a functional description using the standardized instruction. In the second stage, this self-generated description is fed back into the model as a clean input to generate the final embedding. This protocol aligns the semantic content with the model’s native representation space.

### Construction of ground truth gene sets

Gene sets representing known biological pathways were obtained from the Molecular Signatures Database (MSigDB), specifically leveraging four curated collections: Hallmark gene sets, KEGG pathways, Reactome pathways, and Gene Ontology (GO) Biological Process terms. All analyses utilized *Homo sapiens*-specific annotations, which were preprocessed as follows: (i) gene identifiers were standardized to HUGO Gene Nomenclature Committee (HGNC) symbols; (ii) pathways containing fewer than 20 or more than 500 genes were excluded to maintain biological relevance and statistical power. A binary gene-pathway association matrix was constructed, and iterative unsupervised clustering was applied to define functionally coherent and non-overlapping gene modules.

The analytical pipeline of the binary association matrix, implemented in Seurat (v5.1.0)^14^, began with data standardization and dimensionality reduction via principal component analysis (PCA). The top 30 principal components were retained to construct a k-nearest neighbor graph. Identification of major gene modules with distinct functional space was performed through three sequential steps: (1) Initial Module Definition: Unsupervised Louvain clustering^15^ was applied to the binary association matrix to delineate broad functional modules. Module coherence and statistically significant enrichment were confirmed via UMAP visualization and enrichment heatmap analysis. (2) Module Refinement: The top five modules exhibiting the highest biological relevance were subjected to refined Louvain clustering to resolve distinct biological subclusters. These subclusters were validated based on significant enrichment for specific Hallmark gene sets. (3) Ground Truth Curation: The top five most distinct clusters were extracted from the refined subclusters and designated as the ground truth gene sets. Each cluster was annotated with its most significantly enriched Hallmark pathway to define its nomenclature.

All clustering and enrichment analyses were performed using standardized protocols for MSigDB pathway data. Functional enrichment analysis was conducted with clusterProfiler (v4.12.0)^16^ using hypergeometric tests against the Hallmark gene set collection. P values were adjusted for multiple testing using the Benjamini-Hochberg procedure (false discovery rate (FDR) < 0.05). The most significantly enriched Hallmark term was assigned as the cluster label. All analyses were performed in R (v4.4.1) with Seurat (v5.1.0), clusterProfiler (v4.12.0), and msigdbr (v7.5.1).

### Benchmark of gene clustering

Gene embeddings were processed using the Seurat (v5.1.0) pipeline. After loading the embedding matrix, data were normalized and scaled, followed by PCA for dimensionality reduction (30 components retained). K-means clustering (k = 5) was applied directly to the PCA coordinates. For visualization, Uniform Manifold Approximation and Projection (UMAP) was performed using the first 30 principal components. In the resulting two-dimensional plots, genes were color-coded by their ground-truth pathway annotations. UMAP axes were adjusted per embedding type to ensure consistent visualization scales. Clustering performance was quantified by computing the adjusted Rand index (ARI) between the k-means cluster assignments and the ground truth pathway labels using the scikit-learn (v1.5.2) Python package^17^.

### Benchmark of pathway prediction

Pathway prediction was formulated as a supervised multi-class classification task. For each gene embedding set, a random forest classifier (100 decision trees) was trained. Prior to model fitting, embeddings were standardized via z-score normalization and their dimensionality was reduced to 30 components via PCA. Model performance was evaluated using a robust 3×5-fold repeated cross-validation scheme. Given potential class imbalance across pathway categories, performance was primarily assessed using balanced accuracy, which averages the recall across all classes to provide an unbiased estimate.

### Benchmark of Protein–Protein Interaction (PPI) Prediction

A high-confidence human PPI network was constructed by integrating data from three primary databases: STRING (v12.0; interactions with a combined score ≥ 700)^18^, BioGRID (v4.4.248; physical interactions only)^19^, and IntAct (v1.0.4; all interactions)^20^. Data were restricted to Homo sapiens (Taxon ID 9606). Interactions were defined as unique pairs of HGNC gene symbols, and self-interactions were excluded. The final positive dataset, comprising 1,238,984 unique interactions, was formed by taking the union of pairs from these sources.

PPI prediction was framed as a binary classification problem. Individual proteins were represented by their corresponding gene embeddings. For each candidate protein pair, a feature vector was constructed by computing the element-wise product of the two embedding vectors. Features were standardized, and dimensionality was reduced to 30 components via PCA. A Random Forest classifier (100 decision trees) was trained to discriminate true interacting pairs from non-interacting pairs; negative examples were randomly sampled from non-observed protein pairs. Model performance was evaluated using a 3×5-fold repeated cross-validation scheme and reported as the Area Under the Receiver Operating Characteristic curve (AUROC) and the Area Under the Precision-Recall Curve (AUPRC).

### Genome-scale benchmark of human protein-coding genes

Embedding quality was assessed through two complementary approaches: semantic similarity comparison and functional clustering. (1) Semantic Similarity Comparison: The biological consistency of embeddings was evaluated by comparing embedding-based similarity with ontology-based semantic similarity. For each GO Biological Process term, its information content was calculated as the negative binary logarithm of its annotation frequency. A random sample of 1,000 genes was drawn from each embedding set. Pairwise cosine similarity was computed between their embedding vectors and compared to Resnik semantic similarity derived from shared GO annotations. The agreement between the two similarity matrices was assessed using Spearman’s rank correlation, calculated over gene pairs sharing at least one GO annotation. (2) Functional Clustering: After PCA-based dimensionality reduction (30 components), genes were clustered using K-means, with the number of clusters (k) ranging from 30 to 90. For each resulting cluster, enrichment analysis was performed against the GO Biological Process 2025 database. Cluster quality was quantified by calculating the average enrichment score of the top 30 highest-scoring clusters.

## Data availability

All datasets analyzed in this study are publicly available. The ground truth gene sets were constructed from the Molecular Signatures Database (MSigDB). Protein–protein interactions (PPIs) data were derived from the STRING database (v12.0) (https://string-db.org), BioGRID database (v4.4.248) (https://thebiogrid.org/), and IntAct database (v1.0.4) (https://www.ebi.ac.uk/intact/home). The coordinates of human protein-coding genes were obtained from the Ensembl database (GRCh38.p14) using the BioMart tool.

## Code availability

The source code and ground truth dataset are available at https://github.com/hrybiollm/GEbench.

## Acknowledgements

This research work was supported by the Big Data Computing Center of Southeast University.

## Author contributions

PX and YK conceived the concept and designed the study; RH, YK and PX performed the data analyses, prepared the figures, and wrote the paper. YH, WZ, JZ and JL participated in data analysis and manuscript writing. All authors participated in the discussion and interpretation of the results, reviewed and revised the paper.

## Supplementary information

**Additional file 1:** Supplementary Fig. S1.

