## Supplementary material for "Guidance for high-quality functional gene embeddings from large language models": Fig. S1

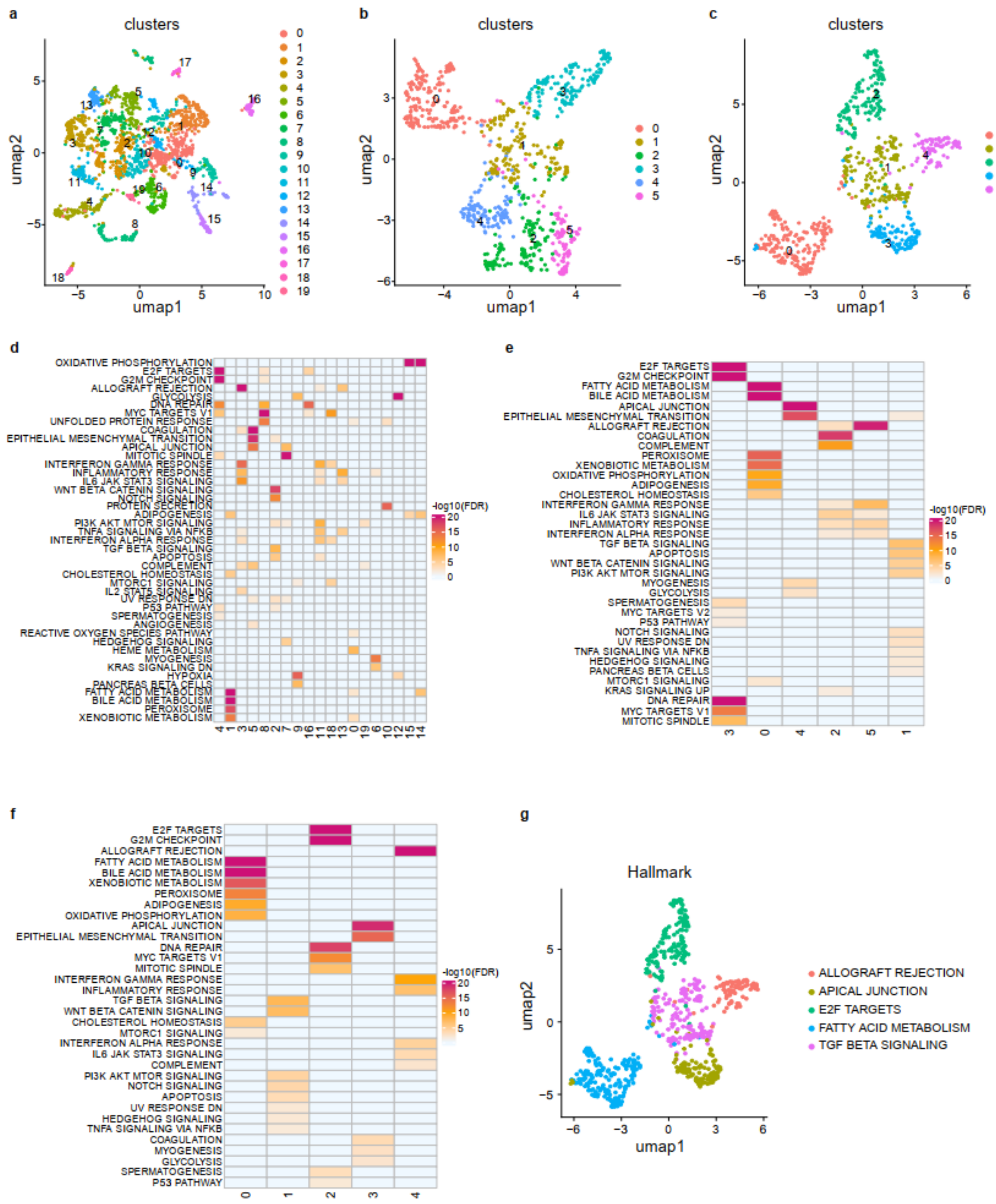


**Supplementary Fig. S1 | Construction of ground-truth functional gene sets via iterative clustering.**

**a,** UMAP visualization of primary pathway modules identified by unsupervised clustering of the binary gene–pathway association matrix constructed from MSigDB collections.

**b,** Refined clustering of the top five primary modules, resolving distinct functional subclusters.

**c,** Selection of five high-confidence, functionally coherent gene clusters from the subclusters in **b** to serve as the GEbench ground truth.

**d–f,** Enrichment heatmaps corresponding to the clustering stages in **a–c**, demonstrating the functional specificity and mutual exclusivity of the selected gene sets.

**g,** Functional annotation of the final five ground-truth gene sets based on their most significantly enriched biological programs.
